# Direct Generation of Images from EEG using Schrödinger Bridge

**DOI:** 10.1101/2025.08.18.670451

**Authors:** Sensho Nobe, Shuntaro Sasai

## Abstract

Real-world data is often noisy, making it challenging to extract true signals Non-invasively recorded neural activities are among the most difficult data, yet its precise signal reconstruction is highly anticipated by communities developing non-invasive brain-machine interfaces. Several noise sources contribute to this challenge, including unrelated neuronal activity, non-brain bioelectricity, attenuation by the skull and scalp, and environmental noises. Additionally, the accumulation of noise varies significantly across subjects and recording sessions, resulting in widely diverging distributions of degraded observations. In this study, we propose modeling the noise accumulation process as a Schrödinger bridge and decoding the true signal by reversing this process. Compared to conventional guided Diffusion approach, our Schrödinger bridge approach effectively models diverse noise processes within a single framework, exhibiting greater robustness to inter-subject variability. Also, our approach doesn’t require pre-aligning brain and image representations, which is an additional compute cost in the conventional approach.

## 1 Introduction

**Figure 1.**
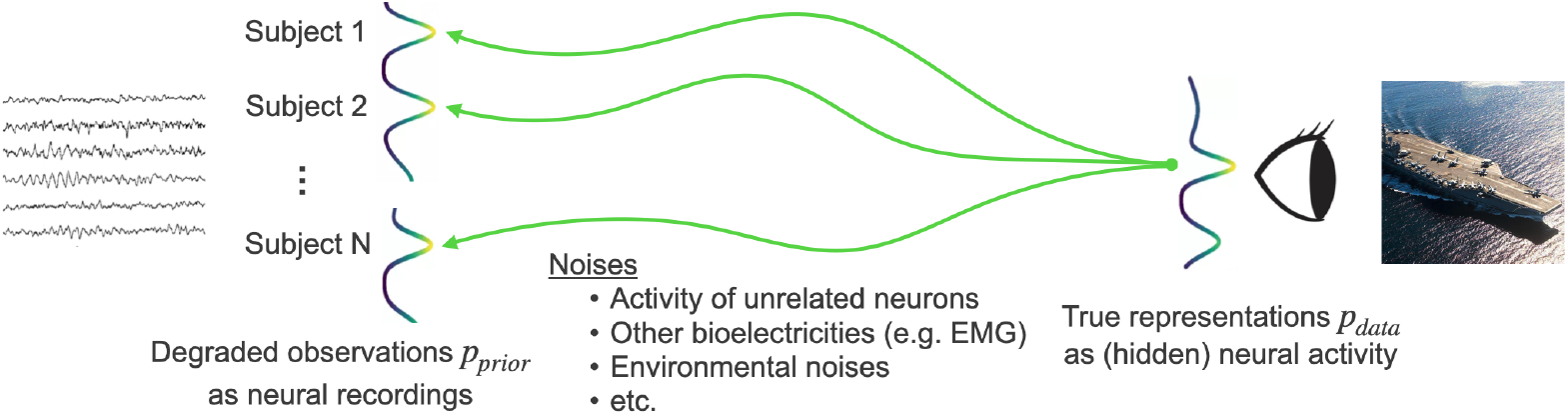
Illustration of noising process of the observed brain activity.

Recent advancements in measurement technologies have significantly improved the accuracy of Brain-Computer Interfaces (BCIs). Techniques like fMRI and MEG provide high spatial resolution across extensive brain areas, enabling the decoding of non-verbal information such as visual images and mental imagery [1, 2, 3, 4]. However, these are large-scale equipment and not portable, which limits their practical application. In contrast, electroencephalography (EEG) offers a more practical solution with less burdensome measurement processes, making it suitable for real-world applications, including those involving healthy individuals. Despite progress in EEG-based decoding of language [5, 6, 7, 8], little progress has been made in decoding visual images. This study aims to develop a framework for the practical application of EEG-based image decoding with an emphasis on mitigating the variabilities of the data.

There are many sources of variability for neural data, such as session-by-session, day-by-day, and subject-by-subject differences, as well as factors such as age and cognitive states, which is a big obstacle in developing stable BCIs [9]. On top of this, EEG signals are notoriously noisy due to various sources of interference before the signal reaches the electrodes, making EEG-based decoding even more challenging. These sources include the activity of numerous unrelated neurons, bioelectrical activities from facial muscles or eye movements, attenuation by the skull and scalp, and environmental noise from recording devices and computers [10]. Previous studies have attempted to address these difficulties by aligning brain embeddings with decoded representations in a CLIP-like manner or by utilizing masked signal modeling of large-scale data [5, 6, 7, 8]. However, training multiple models complicates the pipeline, hindering real-time usage. Furthermore, acquiring large-scale brain activity data for pre-training is not practical. To tackle these issues, our study proposes leveraging the Schrödinger bridge approach, framed within the same context as diffusion models, to directly connect degraded observations with the true signal distribution (Figure 2).

**Figure 2.**
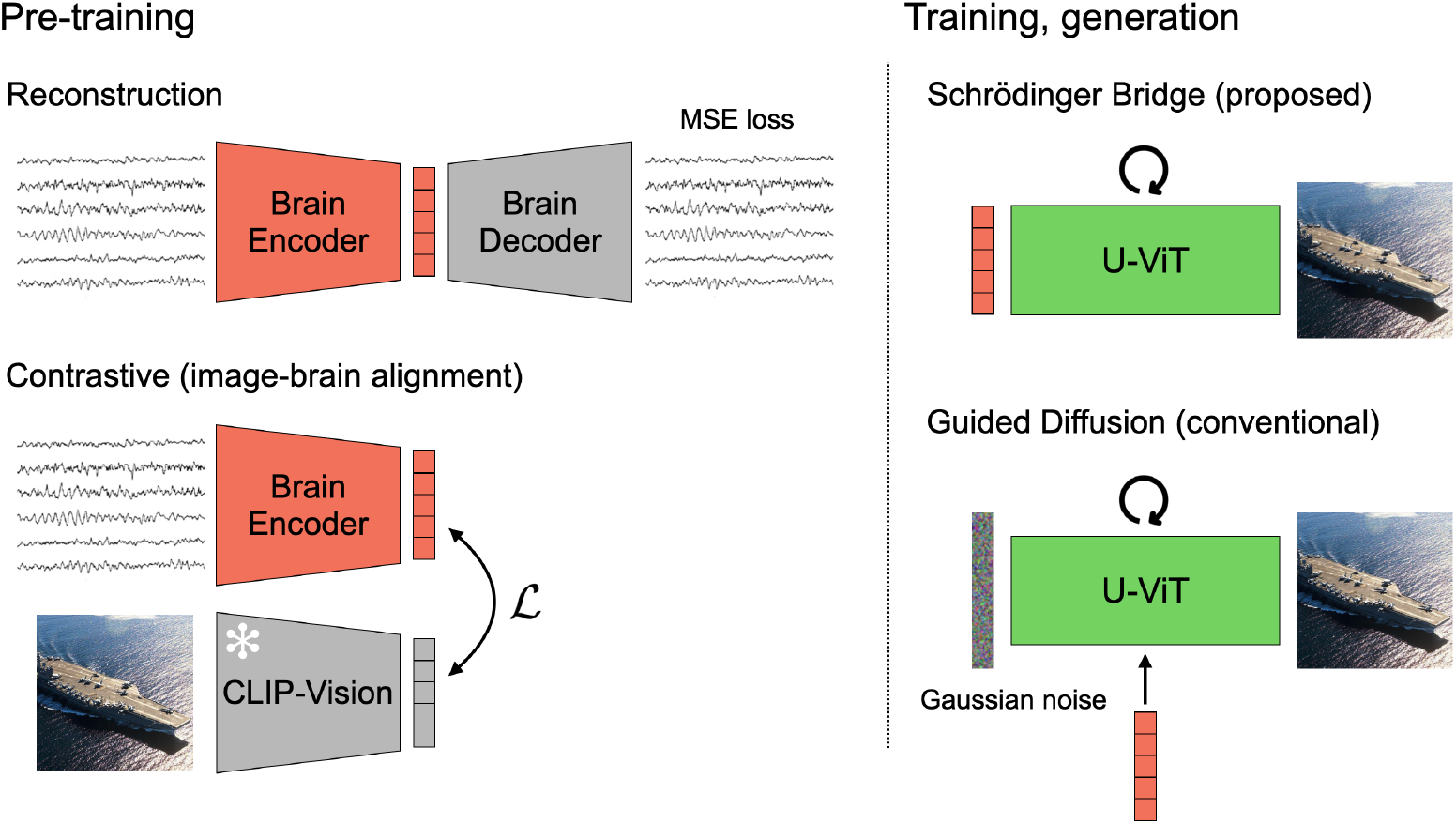
Approaches. Snow mark indicates that the parameters are fixed for the model. We compare reconstruction and contrastive pre-training for both Schrödinger bridge (proposed) and guided Diffusion (conventional) approaches.

## 2 Preliminaries

### 2.1 Diffusion Models and Score-based Generative Models

Diffusion models [11, 12] on a high level are a type of hierarchical variational generative models that considers a process of diffusing the data distribution *p*_data_ to the Gaussian prior distribution *p*_prior_ = 𝒩 (0, *I*) by gradually adding Gaussian noise, and generates samples by modelling to reverse that noising process. The forward noising and backward denoising processes are modelled as a Markov chain with an observed variable **x**_0_ ~ *p*_data_ and latent variables **x**_1:*T*_ where **x**_*T*_ ~ *p*_prior_, the posterior distribution being 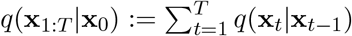. By modelling each step as Gaussian 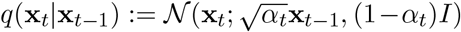 and choosing sufficiently small diffusion steps, we can also model each step of the backward process as Gaussian *p*_*θ*_(**x**_*t−*1_|**x**_*t*_) := N (**x**_*t−*1_; *μ*_*θ*_(**x**_*t*_, *t*), Σ_*θ*_(**x**_*t*_, *t*)). Note that each step of the forward process is a linear Gaussian without a learned parameter, gradually attenuating the true signal. *α*_*t*_ is a time-varying diffusion coefficient that makes the final distribution at time step *T* the standard Gaussian. After the expansion of evidence lower bound of the marginal likelihood, some assumptions such as fixation of the variance to a constant and reparameterization, learning *p*_*θ*_(**x**_*t−*1_|**x**_*t*_) reduces to the training objective ∥***ϵ***_*t*_ − ***ϵ***_*θ*_(**x**_*t*_, *t*)∥^2^, where ***ϵ***_*t*_ is the Gaussian noise added at time step *t*.

The connection from diffusion models to score-based generative models (SGMs) [13] is shown in [14]. Using Tweedie’s Formula [15], they show that the noise prediction objective above can be translated to training a neural network **s**_*θ*_(**x**_*t*_, *t*) that predicts the score function 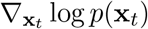. Specifically, the noise and the score function have the following relationship:

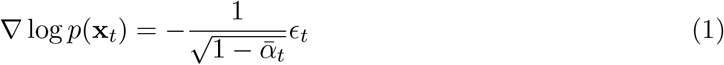

where 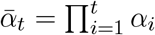.This leads to an interpretation that maximizing the log-likelihood of the data by following the score in the backward process translates to moving in the opposite direction of the noise added in the forward process. Further, if we consider continuous time, the forward and backward processes can be written as the following stochastic differential equations (SDEs):

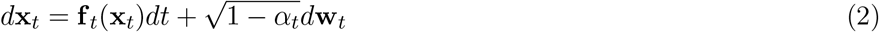

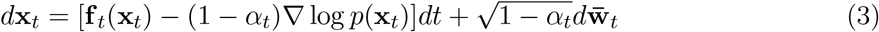

where **f**_*t*_ is the linear drift function, **w**_*t*_ and 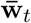 are a standard Wiener process and its time-reversed version, respectively.

### 2.2 Tractable Schrödinger Bridge

Schrödinger bridge [16] can be viewed as an expansion of SGMs where the prior distribution is not limited to the standard Gaussian. In order for the end distribution of the forward process to follow an arbitrary distribution, the non-linear forward drift ∇ log Ψ is added to the forward SDE:

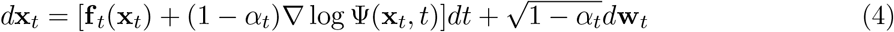

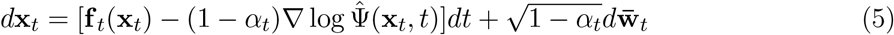

Note that the score function ∇ log *p*(**x**_*t*_) is also replaced with the backward drift 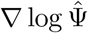, as it is no longer the score function of the forward SDE due to the addition of the non-linear forward drift. Ψ and 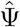 are the energy potentials of the SDEs that satisfy the following coupling constraints:

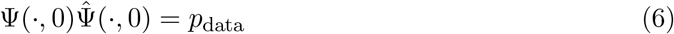

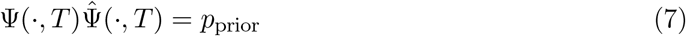

Due to these coupling constraints and inability to reverse the non-linear forward drift, it is not easy to train backward drift 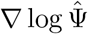 in SGMs framework. [17] address this issue by assuming Dirac delta distribution for data and directly learning a Schrödinger bridge between samples and the prior distribution. While this assumption might decrease generalizability, it breaks one of the coupling constraints and thus training of 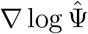 tractable.

## 3 Methods

### 3.1 Dataset and preprocessing

We adopt the ImageNet-EEG paired dataset [18] for testing our method. Brain activity of 6 subjects were recorded while they were presented with one of the 2,000 images chosen from ImageNet, consisted with 50 images each from 40 object classes. Each EEG segment (corresponding to one image) was recorded for 0.5 seconds with 1kHz using a 128-channel headset. We leave more detailed descriptions to the original paper.

As in [18], the first 20ms are discarded from the already epoched raw EEG segments to reduce interference from the previous trials, then they are cropped to 440ms to account for slightly different recording lengths. After applying a notch filter at 50, 100, 150Hz, the segments are centered channel-wise using scikit-learn [19] RobustScaler and values above +/−5 are all clamped. Finally, channel-wise baseline correction is performed using the mean of the first 20% for each sample. From the default train, validation and test splits of the dataset, validation set is included to the train set for data amount. We run both brain encoder pre-training and generation models training with this split, meaning that the whole pipeline sees unknown samples during evaluation.

### 3.2 Brain encoder pre-training

We test two types of pre-training for our brain encoder, where the brain embeddings are aligned to the image embeddings in the second one and not in the first one. The first pre-training is a plain reconstruction task, where a simple decoder of blocks with upsampling, 1d-convolution, layer normalization and GELU [20] activation is added after the brain encoder and trained with mean squared error (MSE) loss. The second pre-training is CLIP [21]-like contrastive one, where the task for the brain encoder is to to maximize the cosine similarity of brain embeddings to CLIP-Vision CLS tokens R^768^ of their paired images while minimizing that to other CLIP-Vision embeddings within batches. Because the cosine similarity only considers angles between vectors, it would be better to match the Euclidean distances to further use the embeddings for image generation. As in [4], we place an additional MSE head and take the combined objective with weighting *λ* = 0.75:

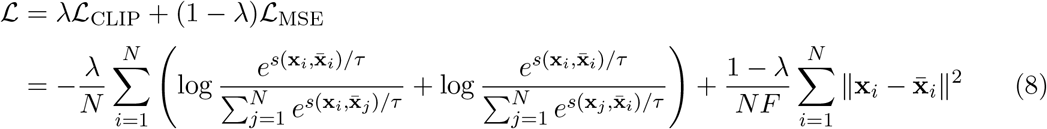

where *s* is the cosine similarity, *τ* is the learnable temperature,**x**_*i*_ is the output embedding of brain encoder, 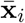 is an CLIP-Vision CLS token to be aligned. Outputs from the MSE head are used for training the generative models.

As the backbone of our brain encoder, we modify the architecture in [4] so that the dilated convolution blocks are replaced with Conformer [22] blocks with an absolute sinusoidal positional encoding at the beginning (see Appendix A.2 for brain encoder backbone selection). Additionally, the dimensionality of the MSE head is increased to ℝ^4096^ to match the Stable Diffusion [23] latent ℝ^4*×*32*×*32^, where we run our noising and denoising processes. CLIP-Vision tokens as the target of the MSE objective are also mapped to ℝ^4096^ with a simple two-layer MLP with layer normalization and GELU activation in the middle. For the reconstruction task, bottleneck dimensionality of the autoencoder is set to ℝ^4096^. As in [6, 4], we introduce subject-specific 1×1 convolution layers that are trained specifically for each subject to account for inter-subject variability. However, Figure 3 shows that the subject layers do not completely absorb the variability, which motivates us for a method robust to these domain shifts.

**Figure 3.**
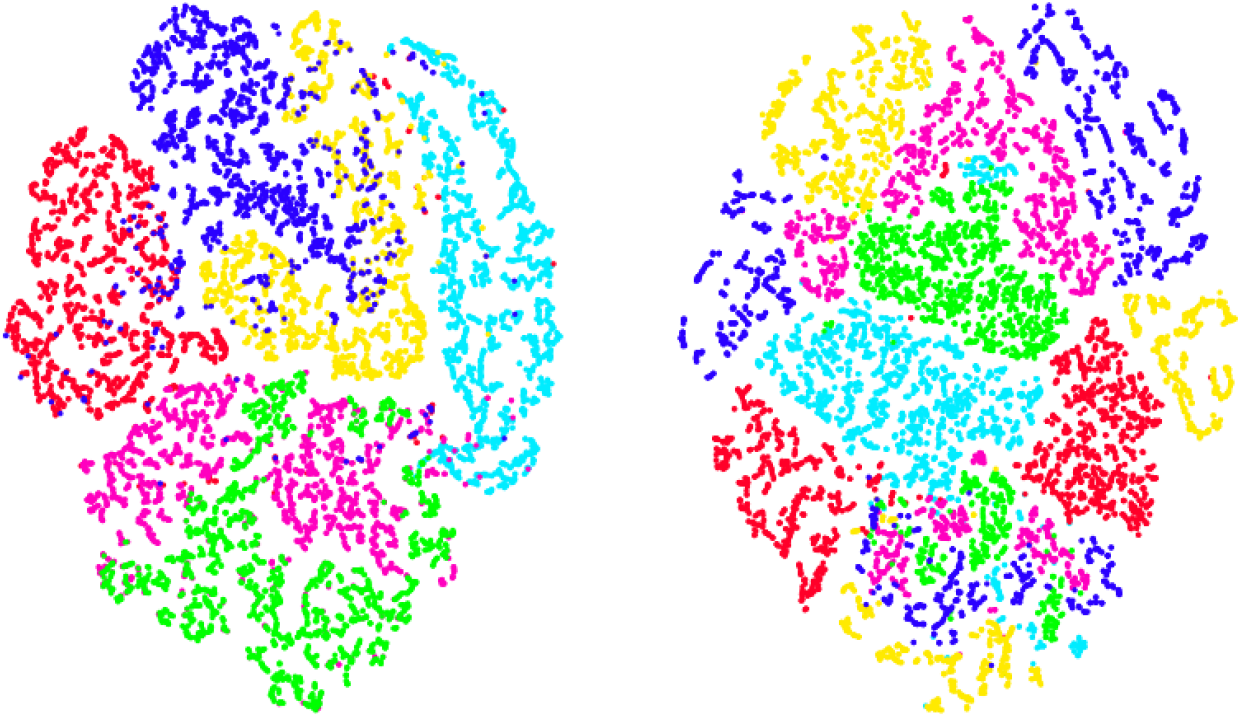
Brain encoder embeddings from the reconstruction task (left) and contrastive task (right) are reduced from ℝ^4096^ to ℝ^2^ using t-SNE with perplexity=10. Latents are colored by 6 subjects. One can see that the subject layers do not completely absorb the between subject variability.

### 3.3 Schrödinger bridge and conditional diffusion training

We parameterize our score prediction network ***ϵ***_*θ*_(**x**_*t*_, *t*) with U-ViT [24]. U-ViT is a recently proposed ViT-based backbone for SGMs with some architectural features in common with U-Net [25], such as long skip connections. U-ViT accepts concatenation of image patches, time, and condition as a sequence of tokens. Originally ℝ^4*×*32*×*32^ Stable Diffusion latents are patched to size 2 and concatenated with time tokens before being input to U-ViT. For conditional diffusion training, we also concatenate a brain encoder embedding as a condition token, which is further mapped linearly to fit in the U-ViT latent. This condition token is replaced with an empty token with probability of 0.15 to jointly train an unconditional model for later guidance sampling.

For Schrödinger bridge training, we sample our noised samples**x**_*t*_ from the discretized version of the posterior discussed in [17]:

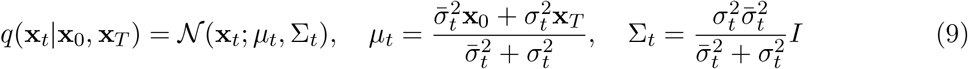

Where 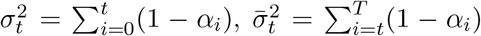, and**x**_0_ and**x**_*T*_ are the image embedding and brain embedding, respectively. As modelling the linearity can be absorbed to the non-linear drift term in the SDEs, we can drop**f** := 0 and then the training objective becomes ∥(**x**_*t*_ −**x**_0_)*/σ*_*t*_ −***ϵ***_*θ*_(**x**_*t*_, *t*)∥^2^ [17]. See Appendix A.3 for more detailed training settings.

### 3.4 Sampling

For guided diffusion, normal DDPM [12] sampling with guidance weight 0.7 is performed. For Schrödinger bridge, DDPM sampling starting from brain encoder embedding**x**_*T*_ is run following the posterior 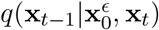, where 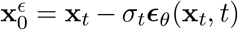. Finally, the sampled embeddings**x**_0_ are fed into the Stable Diffusion decoder to generate the final images for both training.

### 3.5 Evaluation

As high-level evaluation metrics, we report Fréchet inception distance (FID) [26] and the cosine similarity calculated in the CLIP-Vision embedding space (CLIP). For low-level evaluation metric, we report Structural Similarity Index Metric (SSIM).

## 4 Results

Figure 4 shows the images generated from brain activity by our framework. Based on the cosine similarity of CLIP-Vision embeddings between the ground truth and generated images, it displays the most similar 12 images (Best), the 6 images around the median (Average), and the most dissimilar 12 images (Worst). Table 1 shows the evaluation metrics of our approach compared to those of conventional guided diffusion approach. Schrödinger bridge trained with embeddings from reconstruction pre-training shows competitive performances to the diffusion model trained and guided with those from contrastive pre-training, while the diffusion model significantly decreases its performances when the embeddings are not aligned to images beforehand (reconstruction). Interestingly, Schrödinger bridge shows slightly better evaluation results when the embeddings are not aligned, possibly due to the significant difference of the CLIP-Vision representation from that of Stable Diffusion latent. Schrödinger bridge with reconstruction pre-training shows significantly better FID score while showing slightly lower CLIP value than guided diffusion. This might reflect the fact that the samples generated by Schrödinger bridge are much closer to training samples as they are modelled as a Dirac delta distribution, thus having more details than the samples from guided diffusion.

**Table 1:** Evaluation metrics of images generated from Schrödinger bridge and conventional guided diffusion.

| Generation | Pre-training | FID ↓ | CLIP ↑ | SSIM ↑ |
| --- | --- | --- | --- | --- |
| Schrödinger bridge | reconstruction | 29.783 | 0.529 | 0.156 |
|  | contrastive | 35.524 | 0.502 | 0.154 |
| Guided diffusion | reconstruction | 192.701 | 0.437 | 0.214 |
|  | contrastive | 96.592 | 0.568 | 0.144 |

**Figure 4.**
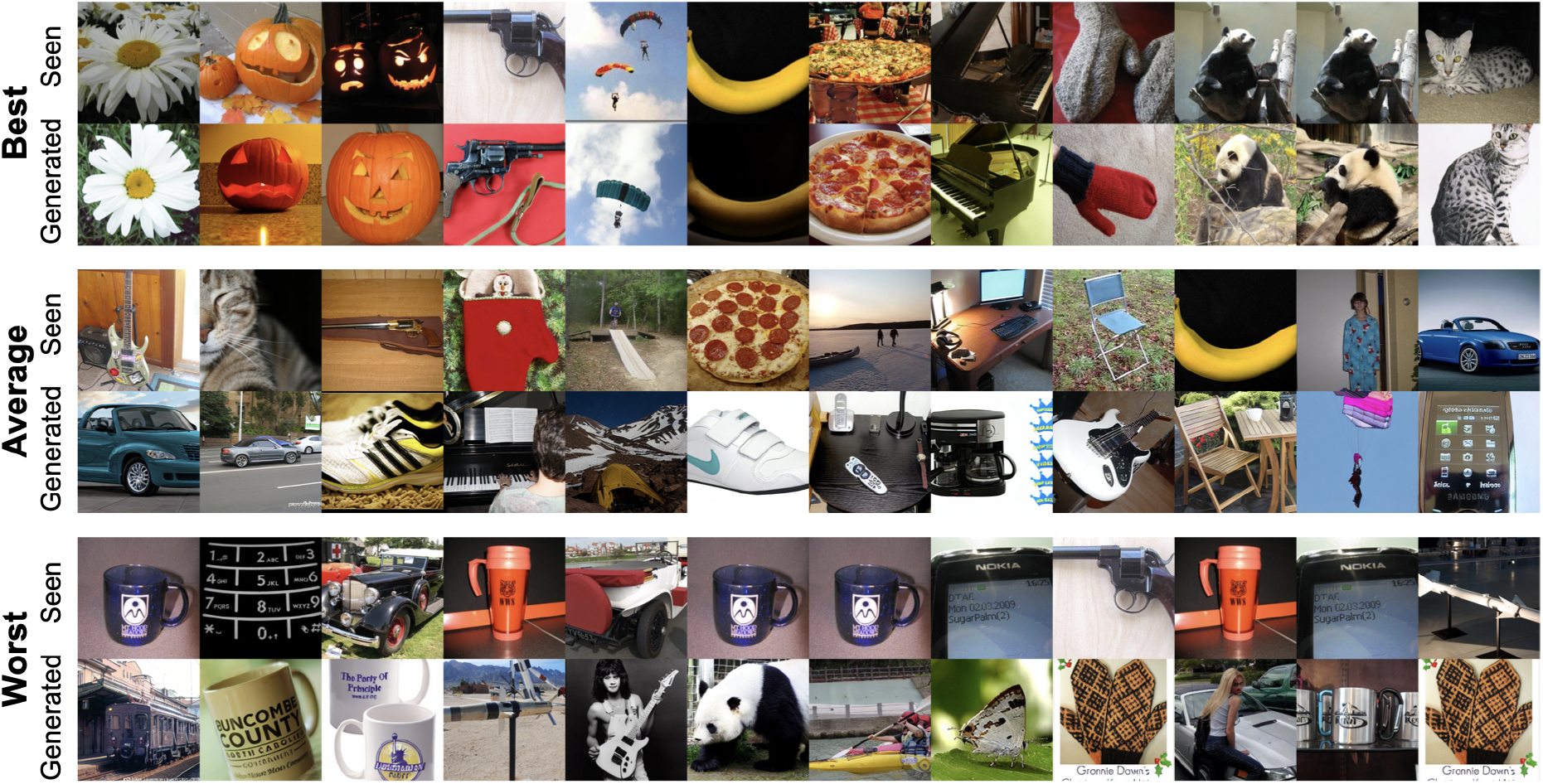
Example images generated from Schrödinger bridge with the brain encoder pre-trained with reconstruction. We display top, middle and worst 12 examples of the CLIP evaluation value. Note that there are duplicate ground truths as there are multiple subjects.

To visualize the confusion matrix of Schrödinger bridge, cosine similarities between the ground truth (seen) and generated images in the CLIP-Vision latent space are calculated and averaged over the 40 object classes (Figure 5). There is a clear trend that animal classes are easier to be mapped from the degraded observations (e.g. giant_panda, African_elephant, Egyptian_cat), especially when the embeddings are aligned by the contrastive pre-training. The possible reason is that the representations of animal images are substantially distinct and thus disentangled in the CLIP space. Additionally, it is probable that this distinctiveness generates unique patterns of brain activity, which makes decoding of those images easier.

**Figure 5.**
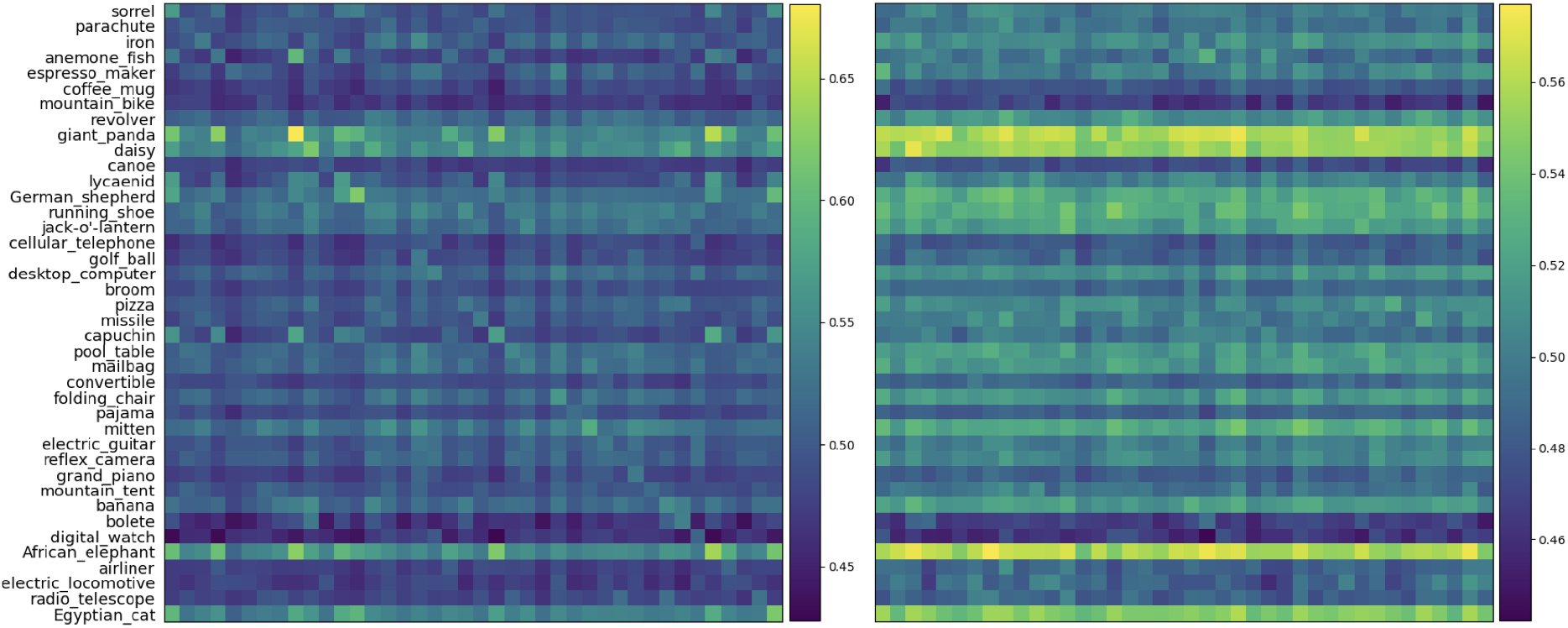
Cosine similarities between ground truth (seen) and generated images in CLIP-Vision latent space averaged over the 40 object classes, for reconstruction (left) and contrastive (right) pre-training.

**Figure 6.**
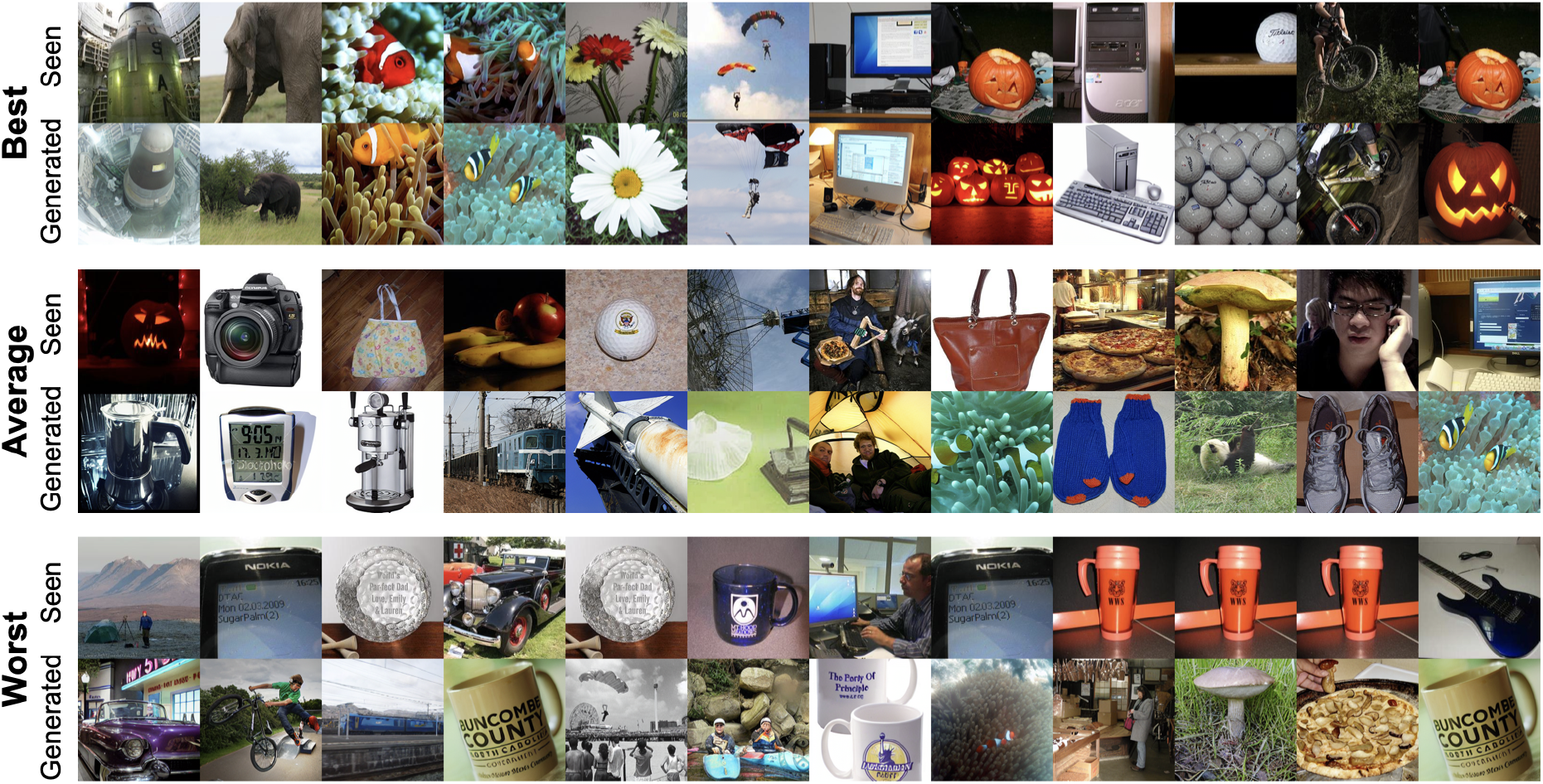
Example images generated from Schrödinger bridge with the brain encoder pre-trained with contrastive task. We display top, middle and worst 12 examples of the CLIP evaluation value. Note that there are duplicate ground truths as there are multiple subjects.

## 5 Limitation

Neural data, especially those obtained by EEG, are substantially variable such that its effective reduction is crucial for utilizing them in applications such as BCIs. This study successfully demonstrated that degraded observations from non-invasive brain recordings can be directly bridged to the true representations we aim to decode. Unlike conventional approaches that condition generative models with embeddings, our method does not require pre-aligning the brain encoder representation and works with the simplest reconstruction pre-training. However, our approach has three main limitations:

1. Limited Generalization: Our method relies on a Schrödinger bridge framework, which is made tractable by assuming a Dirac delta for the data distribution. This assumption limits the generalization ability beyond the training samples.
2. Sampling Duration: The current approach is framed within DDPM sampling, requiring full Markovian steps, which results in a long sampling duration.
3. Generalization to New Subjects: Our method has not been tested for generalization to new subjects.

## 6 Discussion

Previous research on mitigating variabilities of neural data has primarily relied on signal processing techniques. For example, muscle potentials from facial muscles are major confounds in EEG data. A common method involves simultaneously measuring these noise sources, such as EMG, alongside EEG and removing them using linear regression or similar techniques. However, since the true noise-free signal cannot be measured directly, it is challenging to determine the effectiveness of the methods. Furthermore, such methods cannot mitigate significant inter-trial variability and inter-subject variability. Moreover, they rely on the experimenter’s hypothesis about the source of variabilities, which may lead to overlooking the impact of unexpected confounds. In contrast, our approach demonstrates that it is possible to model the process by which noise accumulates without any assumptions, addressing these issues without observations of potential noises. Moreover, while we used EEG data, the proposed method is applicable to any type of data. Thus, it can be broadly applied not only in BCI and brain decoding research but also in general neuroscience research. Future studies should explore applying this method to various data types and tasks to verify whether it can produce stable results surpassing traditional noise reduction techniques.

## Acknowledgements

We thank the creators of the ImageNet-EEG dataset for making their data publicly available. This research was supported by Araya Inc.

## Code availability

Code is available on https://github.com/SeanNobel/sb-image-decoding/.

## A Appendix

### A.1 Generation results with contrastive pre-training

### A.2 Brain encoder model selection and hyperparameters

We follow basic settings of our brain encoder to [4]: two (Conformer or convolutional) blocks, both the spatial attention and subject layers [6], EEG temporal aggregation by affine projection. We run a model selection experiment for top-1 zero-shot classification accuracy, with Adam optimizer of learning rate 3 × 10^*−*4^, batch size of 128, patience of 20 epochs. As for Conformer blocks, we set the number of attention heads to 4, and the depthwise convolution kernel size to 31. The results are listed in Table 2. The experiment is done using one NVIDIA GeForce RTX 3090 GPU, with one training run taking 10 min to 1 hour, depending on early stopping.

**Table 2:** Brain encoder model selection. Top-1 accuracy is calculated within batches, meaning chance-level is 1 / 128 ≃ 0.0078

| Blocks | Position embedding | Test top-1 accuracy |
| --- | --- | --- |
| Conformer | sinusoidal | 0.158 |
| Conformer | learned | 0.141 |
| Dilated conv | - | 0.086 |

### A.3 U-ViT hyperparameters and training

Same configurations of U-ViT hyperparameters are used for Schrödinger bridge and guided diffusion training: latent dimension of 1024, block number of 20, attention head number of 16, feedforward dimension of 4096, and patch size of 2. Following Stable Diffusion, we use a quadratic noise schedule with *β*_*t*_(= 1 − *α*_*t*_) starting from 0.00085 and ending in 0.012 for guided diffusion. For Schrödinger bridge, we use a symmetric scheduling where *β*_*t*_ value is maximized to 0.0057 in the middle of the DDPM steps and minimized to 0.0001 at both ends. DDPM steps are set to *T* = 1000 for both. We train U-ViT for 100,000 steps with mixed precision (16-bit) using 4 NVIDIA RTX A5000 GPUs, with one run taking approximately 5 days.

## References

[1] Furkan Ozcelik and Rufin VanRullen. “Natural scene reconstruction from fMRI signals using generative latent diffusion”. In: Scientific Reports 13.1 (Sept. 2023), p. 15666. ISSN: 2045-2322.

[2] Sikun Lin, Thomas Sprague, and Ambuj K Singh. “Mind Reader: Reconstructing complex images from brain activities”. In: Advances in Neural Information Processing Systems. Ed. by S. Koyejo et al. Vol. 35. Curran Associates, Inc., 2022, pp. 29624–29636.

[3] Paul Scotti et al. “Reconstructing the Mind’s Eye: fMRI-to-Image with Contrastive Learning and Diffusion Priors”. In: Advances in Neural Information Processing Systems. Ed. by A. Oh et al. Vol. 36. Curran Associates, Inc., 2023, pp. 24705–24728.

[4] Yohann Benchetrit, Hubert Banville, and Jean-Remi King. “Brain decoding: toward real-time reconstruction of visual perception”. In: The Twelfth International Conference on Learning Representations. 2024.

[5] Young-Eun Lee et al. “Towards voice reconstruction from EEG during imagined speech”. In: Proceedings of the AAAI Conference on Artificial Intelligence. Vol. 37. 5. 2023, pp. 6030–6038.

[6] Alexandre Défossez et al. “Decoding speech perception from non-invasive brain recordings”. In: Nature Machine Intelligence 5.10 (2023), pp. 1097–1107.

[7] Zhenhailong Wang and Heng Ji. “Open vocabulary electroencephalography-to-text decoding and zero-shot sentiment classification”. In: Proceedings of the AAAI Conference on Artificial Intelligence. Vol. 36. 5. 2022, pp. 5350–5358.

[8] Yiqun Duan et al. “DeWave: Discrete Encoding of EEG Waves for EEG to Text Translation”. In: Thirty-seventh Conference on Neural Information Processing Systems. 2023.

[9] Alan D Degenhart et al. “Stabilization of a brain–computer interface via the alignment of low-dimensional spaces of neural activity”. In: Nature biomedical engineering 4.7 (2020), pp. 672–685.

[10] Saeid Sanei and Jonathon A Chambers. EEG signal processing. John Wiley & Sons, 2013.

[11] Jascha Sohl-Dickstein et al. “Deep Unsupervised Learning using Nonequilibrium Thermodynamics”. In: Proceedings of the 32nd International Conference on Machine Learning. Ed. by Francis Bach and David Blei. Vol. 37. Proceedings of Machine Learning Research. Lille, France: PMLR, July 2015, pp. 2256–2265.

[12] Jonathan Ho, Ajay Jain, and Pieter Abbeel. “Denoising Diffusion Probabilistic Models”. In: Advances in Neural Information Processing Systems. Ed. by H. Larochelle et al. Vol. 33. Curran Associates, Inc., 2020, pp. 6840–6851.

[13] Yang Song et al. Score-Based Generative Modeling through Stochastic Differential Equations. 2021. 2011.13456 [cs.LG].

[14] Calvin Luo. Understanding Diffusion Models: A Unified Perspective. 2022. 2208.11970 [cs.LG].

[15] Bradley Efron. “Tweedie’s Formula and Selection Bias”. In: Journal of the American Statistical Association 106.496 (2011). PMID: 22505788, pp. 1602–1614.

[16] Christian Léonard. A survey of the Schrödinger problem and some of its connections with optimal transport. 2013. 1308.0215 [math.PR].

[17] Guan-Horng Liu et al. “I^2^SB: Image-to-Image Schrödinger Bridge”. In: Proceedings of the 40th International Conference on Machine Learning. Ed. by Andreas Krause et al. Vol. 202. Proceedings of Machine Learning Research. PMLR, 23–29 Jul 2023, pp. 22042–22062.

[18] MM ’17: Proceedings of the 25th ACM international conference on Multimedia. New York, NY, USA: Association for Computing Machinery, 2017. ISBN: 9781450349062.

[19] Fabian Pedregosa et al. “Scikit-learn: Machine learning in Python”. In: Journal of machine learning research 12.Oct (2011), pp. 2825–2830.

[20] Dan Hendrycks and Kevin Gimpel. Gaussian Error Linear Units (GELUs). 2023. 1606.08415 [cs.LG].

[21] Alec Radford et al. “Learning Transferable Visual Models From Natural Language Supervision”. In: Proceedings of the 38th International Conference on Machine Learning. Ed. by Marina Meila and Tong Zhang. Vol. 139. Proceedings of Machine Learning Research. PMLR, 18–24 Jul 2021, pp. 8748–8763.

[22] Anmol Gulati et al. Conformer: Convolution-augmented Transformer for Speech Recognition. 2020. 2005.08100 [eess.AS].

[23] Robin Rombach et al. “High-resolution image synthesis with latent diffusion models”. In: arXiv [cs.CV] (Dec. 2021).

[24] Fan Bao et al. “All Are Worth Words: A ViT Backbone for Diffusion Models”. In: Proceedings of the IEEE/CVF Conference on Computer Vision and Pattern Recognition (CVPR). June 2023, pp. 22669–22679.

[25] Olaf Ronneberger, Philipp Fischer, and Thomas Brox. U-Net: Convolutional Networks for Biomedical Image Segmentation. 2015. 1505.04597 [cs.CV].

[26] Martin Heusel et al. “GANs trained by a two time-scale update rule converge to a local Nash equilibrium”. In: arXiv [cs.LG] (June 2017).

